# Skewness in bee and flower phenological distributions

**DOI:** 10.1101/2022.05.17.492369

**Authors:** Michael Stemkovski, Rachel G. Dickson, Sean R. Griffin, Brian D. Inouye, David W. Inouye, Gabriella L. Pardee, Nora Underwood, Rebecca E. Irwin

## Abstract

Phenological distributions are characterized by their central tendency, breadth, and shape, and all three determine the extent to which interacting species overlap in time. Pollination mutualisms rely on temporal co-occurrence of pollinators and their floral resources, and while much work has been done to characterize the shapes of flower phenological distributions, similar studies including pollinators are lacking. Here, we provide the first broad assessment of skewness, a component of distribution shape, for a bee community. We compare skewness in bees to that in flowers, related bee and flower skewness to other properties of their phenology, and quantify the potential consequences of differences in skewness between bees and flowers. Both bee and flower phenologies tend to be right-skewed, with a more exaggerated asymmetry in bees. Early-season species tend to be the most skewed, and this relationship is also stronger in bees than in flowers. Based on a simulation experiment, differences in bee and flower skewness could account for up to 14% of pair-wise overlap differences. Given the potential for interaction loss, we argue that difference in skewness of interacting species is an under-appreciated property of phenological change.

## Introduction

Timings of seasonal life-history events (phenology), are often characterized by single points in time (e.g., first-appearance date), but in reality these events are typically distributed processes (Carter et al., 2018). Species phenological distributions are characterized by their central tendency, their breadth, and their shape (e.g., mean, standard deviation, and skewness) (Rathcke et al., 1985). Ecological interactions usually require temporal co-occurrence, where the population performance of any species is dependent on phenological overlap with resource availability. In the case of pollination and other mutualistic interactions, interacting species benefit from maximizing temporal overlap with one another, while dealing with the fitness costs of changing their phenology (Visser et al., 2012). The degree of overlap between interacting species is determined by the mean, breadth, and shape of both species’ phenological distributions (Fig. 1), with differences in any one of the three properties being enough to reduce overlap. While the shape of phenological distributions has been recognized as an important component of species interactions (Thomson, 1980), studies of phenological match/mismatch in plants and pollinators have focused primarily on how first or mean dates of seasonal activity shift in response to varying cues (Inouye et al., 2019) and how the temporal breadth of their activity stretches or contracts. The importance of skewness differences in determining mismatch in pollination interactions remains unclear. This is in part because there has not been a systemic analysis of skewness in phenological distributions across many species of pollinators and flowering plants together.

**Figure 1.**
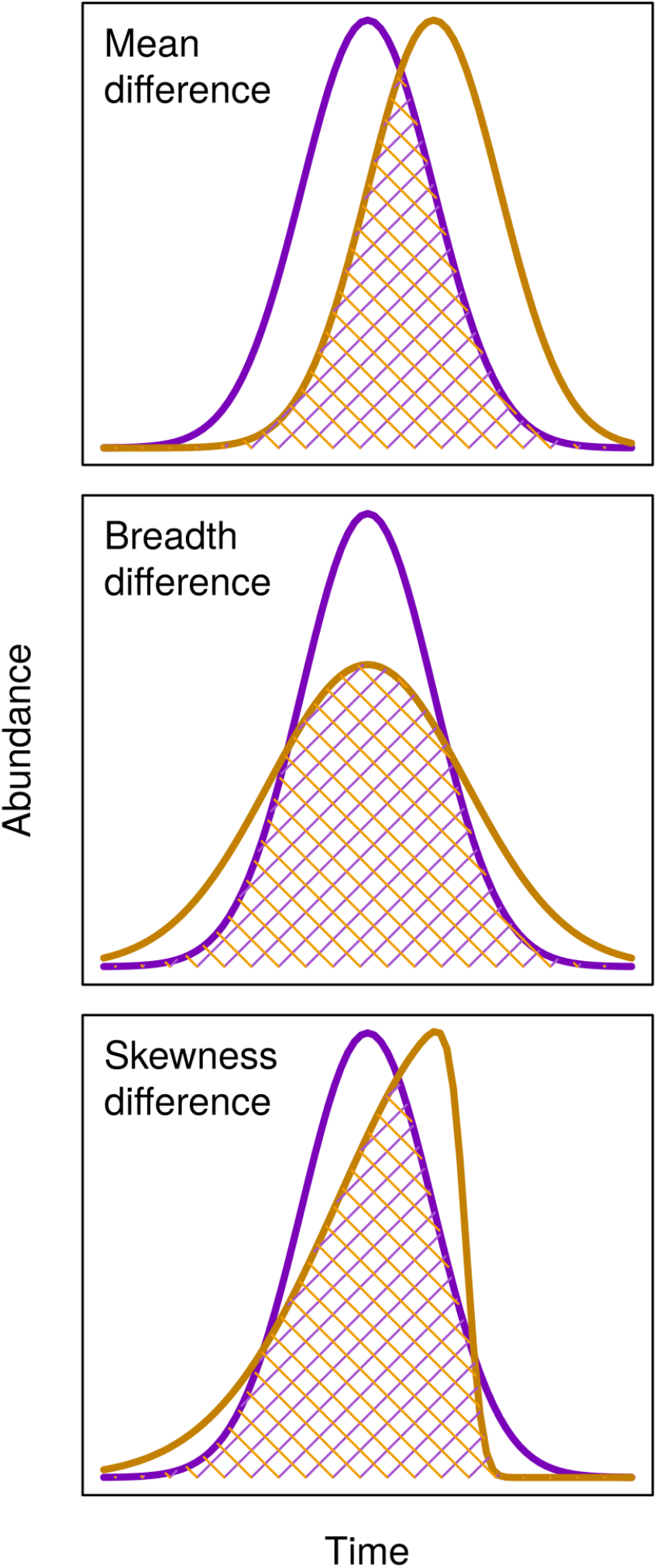
Conceptual diagram of the causes of phenological mismatch. Differences in the phenological mean timing (top panel), breadth (center panel), and skewness (bottom panel) of species determine the extent to which interacting species overlap in time. The purple and yellow curves represent phenological distributions of two species, and the hatched areas are times of phenological overlap.

Flower phenological distributions are often right-skewed, with a long, trailing tail after the peak of flowering in prairie (Rabinowitz et al., 1981) and montane (Thomson, 1980) ecosystems. Within populations, the degree of skewness can vary among years (Blionis et al., 2001; Forrest et al., 2010). Phenological distribution shape may also be affected by local resource competition that reduces plant size, which is in turn correlated with floral phenology skewness (Schmitt, 1983). It has also been suggested that the right-skewness of flowering phenologies may be a product of selective pressure for early flowering (Forrest et al., 2010) and that recent climate change has affected the shapes of flower phenological distributions (CaraDonna et al., 2014). While the typical patterns of flower phenological skewness are well understood, we do not know if these patterns are also similar for pollinators. There is reason to suspect that phenological sensitivity differs between plants and pollinators and that the seasonal onset and end of activity in pollinators (such as bees) shift at different rates (Stemkovski et al., 2020). Thus, a community-wide assessment to compare bee and flower phenological skewness is warranted.

In this study, we quantified phenological skewness for multiple bee and flower species within a montane community. We determined the relative prevalence of right-skewed, left-skewed, and symmetrical distributions, and examined the differences between bees and flowers in how skewness relates to other properties of their phenological distributions. *A priori*, we predicted that bee phenologies would be similarly skewed to those of flowers, but that bees active in the early season would have phenologies more strongly right-skewed than that in the late season due to a hard limit on activity before snowmelt in the study system. Lastly, we performed a simulation study to gain perspective on the potential consequences that variation in skewness in this community may have on phenological overlap between bees and flowers.

## Methods

### Data sources

We used flower phenology data from a long-term monitoring program spanning from 1974 to 2019 at the Rocky Mountain Biological Laboratory (RMBL) in Gothic, Colorado, USA, a mountainous location with a short summer growing season set ~2890 m above sea level (CaraDonna et al., 2014; Inouye, 2008). The total number of flowers was counted approx. three times per week for the extent of the growing season for all flowering species in 4 m^2^ fixed plots. This dataset includes mainly long-lived perennial forb species. Further details on this program are reported by CaraDonna et al. (2014) and Inouye (2008), and all data are available through Open Science Framework (https://osf.io/hy59v/). For our analysis, we included 35 flower plots aggregated into 8 sites by proximity to agree with the spatial scale of the bee phenology sites. We obtained bee phenology data from a companion study to the flower phenology project which tracked bee abundance from 2009 to 2020 at the RMBL (Ogilvie et al., 2017; Stemkovski et al., 2020). Bee abundance was measured using pan traps approx. every two weeks across the growing season at 18 sites spaced across an elevation gradient (Gezon et al., 2015). Because pan traps are biased toward collecting smaller-bodied bees, hand netting was used for bumble bees (*Bombus* spp.). Additional details on bee data collection and all data are available through Open Science Framework (https://osf.io/kmxyn/). All data processing steps and analyses for this study can be viewed and reproduced using code available online (https://github.com/stemkov/pheno_skew).

### Data processing

We formatted and standardized flower and bee data to make them directly comparable. For all data, we excluded records with uncertain identifications and those that were identified only to genus. We excluded all grass and sedge species, but included shrubs. The bee abundance data were derived from multiple pan traps or netters, so we aggregated flower and bee counts across plots/traps/netters per site. For the bee data, we included only female bees because female specimen identifications were more fully resolved and because combining females and males could lead to inaccurate estimates of skewness when the two sexes have different phenological patterns (as in social species). We distinguished queen and worker castes of bumble bees (*Bombus spp*.) to avoid biasing skewness estimation by confounding an early-season queen peak in abundance and a later peak in worker abundance. Because bumble bees were sampled explicitly by netting and due to difficulties of combining sampling effort between netting and pan traps, we excluded pan-trapped *Bombus* and net-trapped non-*Bombus* bees. To ensure that we only included sampling periods that consistently captured representative samples of abundance, we excluded sampling days when traps were deployed for less than three hours and excluded netting days with less than one hour of effort (excluding 17 of 778 trap sampling days and 21 of 809 net sampling days). Lastly, to ensure adequate sample size and robust skewness estimation, we only considered time-series with at least 10 individual bee records and at least 100 flower records. Thus, we excluded 1,932 of 18,710 bee records (10.3%) and 126,659 of 3,943,796 flower records (3.2%).

### Skewness calculation and predictors

We calculated skewness as the Fisher-Pearson standardized third-moment coefficient of skewness (*g_1_*), as implemented in the *moments* R-package (Komsta et al., 2015), for each site/year/species phenological abundance distribution (i.e., frequencies of bee and flower abundance by date). We tested whether skewness was different from zero (corresponding to a symmetrical distribution) using D’Agostino’s K^2^ test (D’Agostino, 1970).

In order to examine whether and how skewness in bees and flowers was related to other properties of their phenological abundance distributions, we calculated the means and standard deviations of each distribution. To test whether the phenological position of species (how early or late they are active in a season) predicted their skewness, we modeled skewness as a linear function of phenological mean interacting with guild (flowers vs. bees). We note that, statistically, means are shifted by skewness, so the two are necessarily linked to a certain extent. To test whether species with longer active seasons tended to be more skewed, we modeled absolute skewness as a linear function of distributions’ standard deviation interacting with guilds.

### Overlap calculation

To demonstrate the potential phenological match/mismatch consequences of skewness differences between bees and flowers, we calculated the maximum possible overlap of distributions with different skews. To do this, we repeatedly generated probability densities of two skew-normal distributions (Azzalini, 2020) with two different skewness parameters and 2000 possible mean and standard deviation parameter combinations each. For each mean and standard deviation value, we then calculated the overlap coefficient (Inman et al., 1989) by integrating to find the area encompassed by both probability density curves (as illustrated in Fig. 1), and recorded the largest of the resulting overlap coefficients. We repeated this procedure for every pairwise combination of 50 skewness values sequenced evenly between −5 and 5, resulting in 2500 (i.e., 50^2^) total comparisons. In other words, we calculated the largest possible overlap of paired distributions by keeping skewness constant and allowing mean and standard deviation to vary freely. To provide perspective, we calculated the bounds of the middle 95% of skewness values of bees and flowers from the Rocky Mountain dataset and overlayed these onto the simulated overlap estimates.

## Results

We estimated skewness for 3,024 flower time-series and 480 bee time-series. The time-series represented 106 plant species across 82 genera, and 49 bee species across 14 genera. In time-series with sufficient data to calculate skewness, the average flowering period (across all years, sites, and species) was centered on July 10, and bee foraging on July 2. The typical flowering breadth, measured as one standard deviation on either side of center, was 14 days in flowers and 39 days in bees. Flower timeseries were significantly right-skewed (*g_1_* = 0.31, t_3502_ = 20.43, p < 0.01), and bee time-series were also right skewed (*g_1_* = 0.89), significantly more so than flowers (t_3502_ = 13.84, p < 0.01). Viewed individually, 47.6% of flower time-series were significantly right-skewed, only 14% were significantly left-skewed, and 38.5% were not significantly different from symmetrical. Of bee time-series, 48.5% were significantly right-skewed, 9.4% were left-skewed, and 42.1% were symmetrical (Fig. 2). Skewness was somewhat affected by data truncation, though both bees and flower curves were still right-skewed regardless of truncation type (Appendix S1: Section 1).

**Figure 2.**
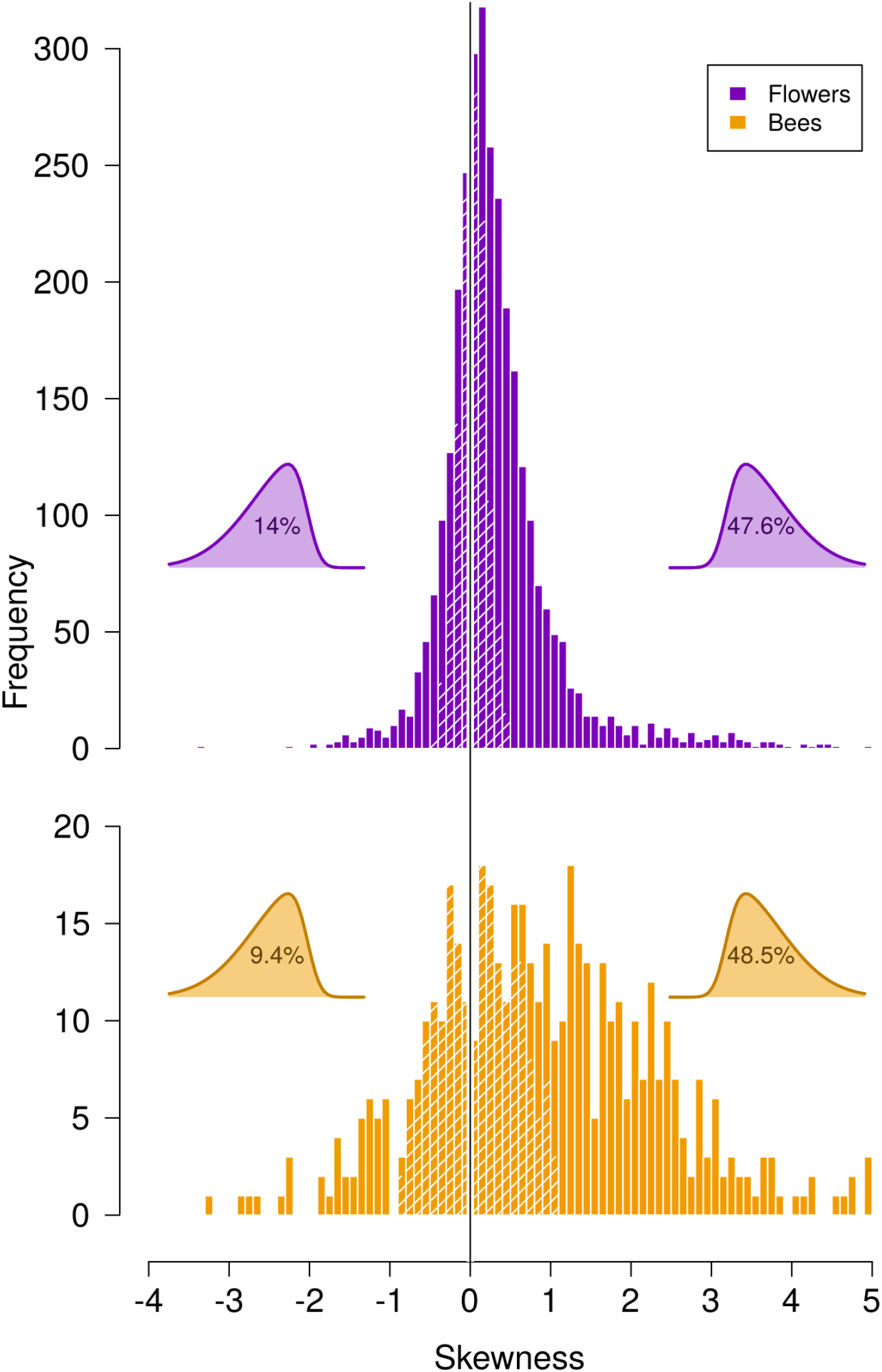
Flower (top panel) and bee (bottom panel) phenological skewness. Both flower and bee species tend to have right-skewed phenological distributions, though there is substantial variation in shape, and most distributions in both groups are not significantly different from symmetrical. Skewed distribution icons give the percent of individual time-series that are significantly left- and right-skewed.

Skewness in plants was significantly predicted by how early or late in the season a species flowered, with early season species being more strongly right-skewed (t_3500_ = −6.77, p < 0.01). This relationship was more pronounced in bees (t_3500_ = −22.8, p < 0.01), with early season bees being right-skewed, and later-season bees being left-skewed (Fig, 3, left panel). Plants with longer flowering periods tended to be more skewed (t_3500_ = 9.32, p < 0.01), while bees with longer active periods tended to be less skewed (t_3500_ = −11.75, p < 0.01) (Fig. 3, right panel). We do not directly compare bee and flower distribution breadth because the frequency of bee data collection inherently discounted species with short active periods. Hypothetical overlap losses between interacting species ranged from 0% at perfectly matched skewness values to 25% for distributions where one has *g_1_* = 5 and the other *g_1_* = −5 (Fig. 4). The maximum overlap loss for the central 95% of bee and flower skewness values found in our datasets was 14% overlap loss. Comprehensive species lists and summary statistics are provided in Appendix S1: Section 3.

**Figure 3.**
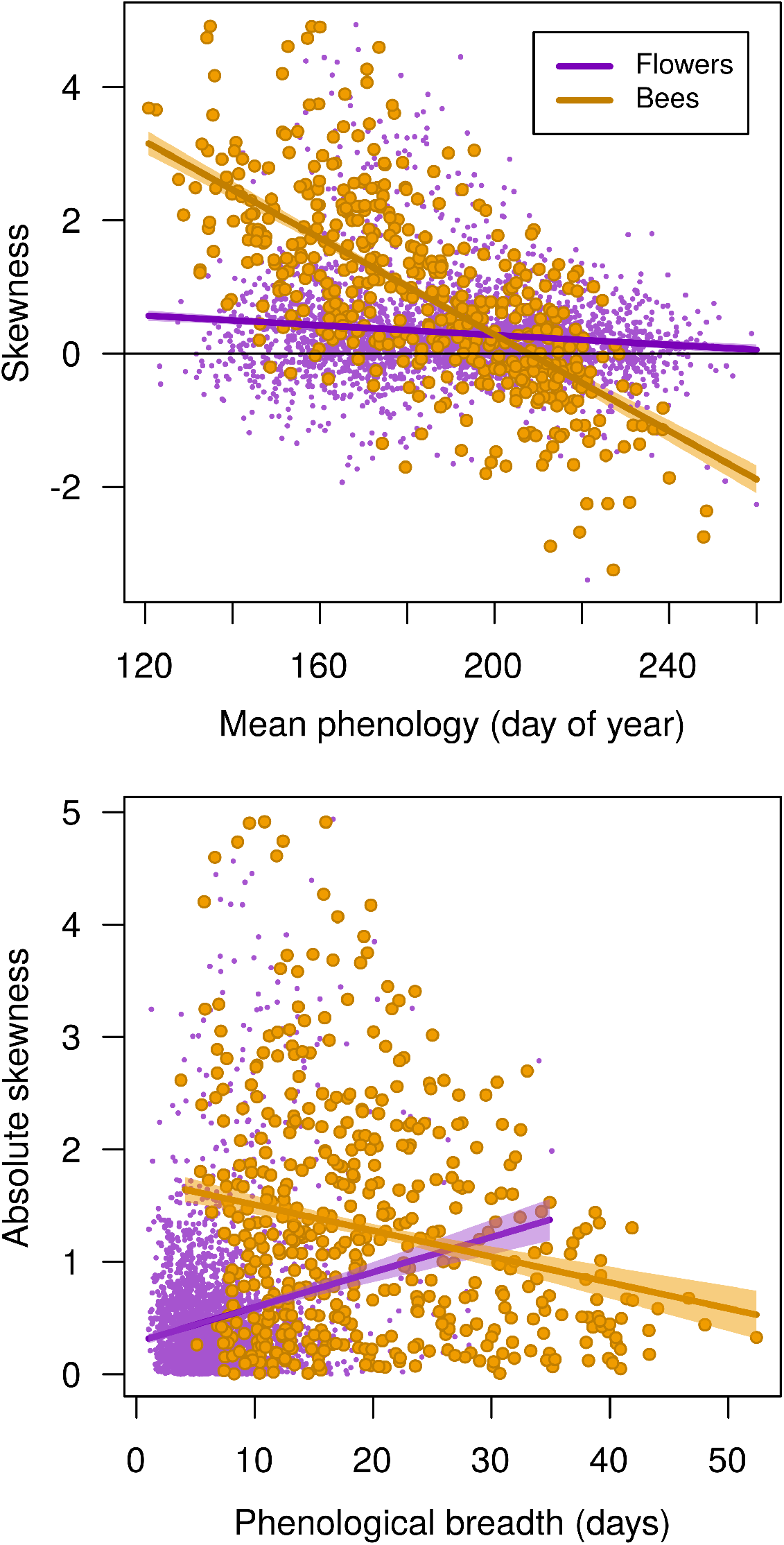
Skewness is predicted by mean and breadth. Early-season bees and flowers tended to be more heavily right-skewed (top panel), thought the effect was more pronounced in bees than in flowers. Flowers with broader phenological distributions tended to be more skewed, while bees with broader phenology tend to be less skewed in either direction (bottom panel).

**Figure 4.**
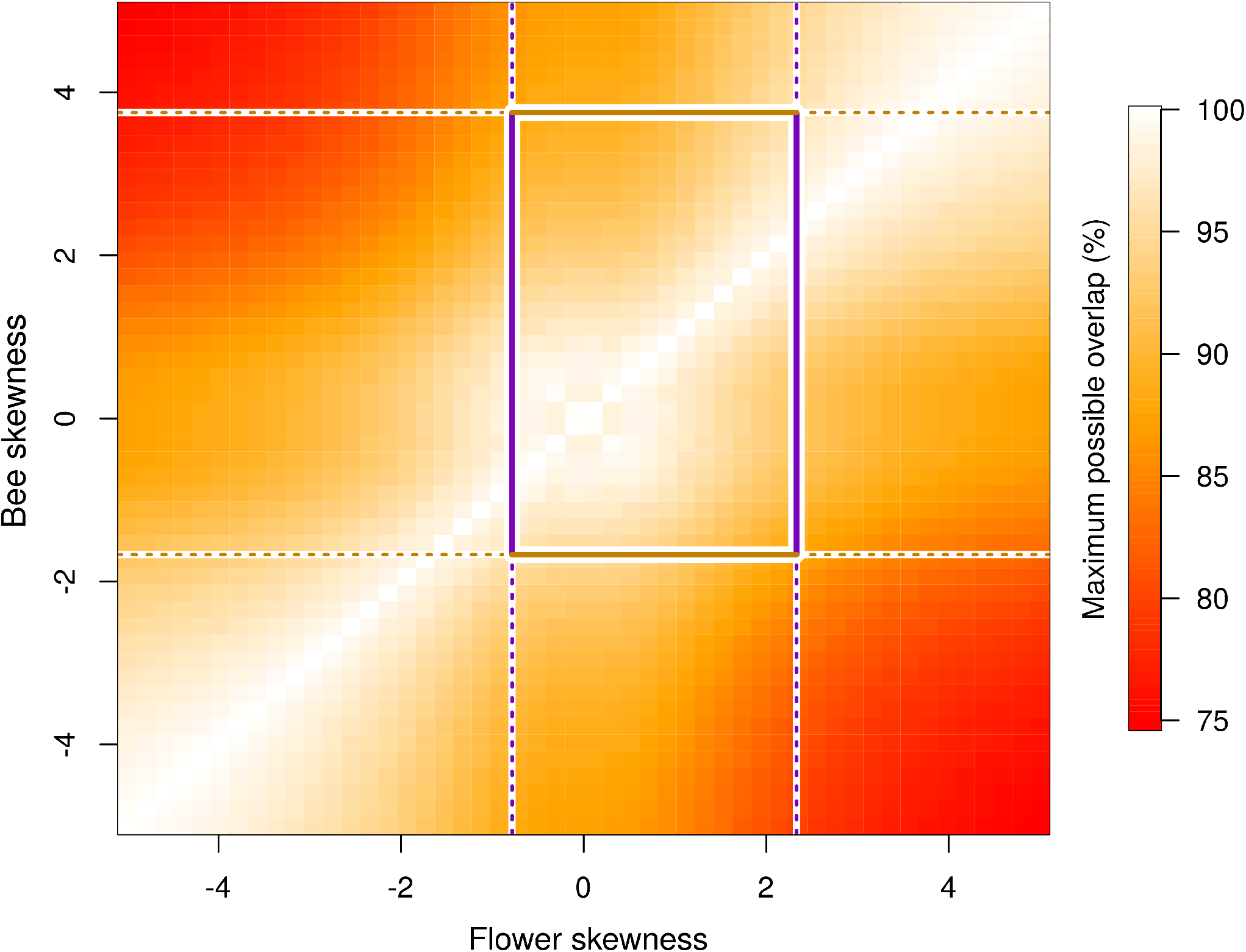
Skewness constrains the degree to which phenological distributions can overlap. The maximum possible overlap of pair-wise interacting species with different hypothetical skewness values is shown as colors, with red depicting the lowest overlap, and white depicting perfect overlap. Bounds for 95% of the actual observed skewness values are shown with purple lines for flowers, and gold lines for bees. The resulting box in the middle of the figure therefore depicts the potential loss of phenological overlap in pair-wise interactions between bees and flowers due to differences in skewness alone, isolated from the effects of mean and breadth changes.

## Discussion

We found that both bee and flower phenological distributions tend to be right-skewed (Fig. 2), suggesting that similar processes are acting on bees and plants to determine the shape of their phenological distributions. Multiple explanations have been offered for skewness in flowering time, including selection for skewed flowering driven by pollinators and resources (Forrest et al., 2010; Thomson, 1980), by-products of intraspecific competition (Schmitt, 1983), and simple geometrical necessity because daily survival probabilities are cumulative and inherently skewed (Blionis et al., 2001). We can identify multiple, distinct scenarios that may lead to skewed phenological distributions by focusing on onset rates (e.g., flower opening, bee emergence) and cessation rates (e.g., floral senescence, bee mortality). First, differences in the intra-annual dispersion of onset and cessation rates may produce phenological skewness. Second, variable phenology among individuals with unequal representation (e.g., different numbers of flowers per individual plant) may produce skewed aggregate distributions even when the component onset and end rates are equally dispersed. Third, onset and/or cessation distributions themselves may be skewed. Further, combinations of these processes may influence the shapes of phenological distributions in complex ways. In the context of our findings, the prevalence of right-skewness in bee phenological distributions suggests that, on average, bees emerge with more synchrony than with which they cease foraging. Drawing conclusions about the processes behind the observed skewness in flower phenology is more difficult due to flower counts being aggregated across individual plants in our data.

While demographic data (i.e., tracking individual plants and insects) beyond what we present here are needed to determine the mechanistic causes of the differences in onset/end variance that produce skewed phenology, some information can be gleaned by comparing skewness with other phenological properties. We found that bees that were active closer to the beginning of the growing season tended to be more right-skewed (Fig. 3). For example, the early season sweat bee *Lasioglossum sedi*, with an average capture date of June 16 across all sites and years, was strongly right-skewed (*g_1_* = 1.94), while the later-season masked bee *Hylaeus annulatus*, with a capture date of August 5 on average, tended to be left-skewed (*g_1_* = −0.27). A similar but weaker pattern was seen in flowers, though flowers did not tend to flip to left-skewness at the end of the season. In the snowy sub-alpine environment of the present study, the onset of activity is strongly limited for flowers (Inouye, 2008) and bees (Stemkovski et al., 2020) by the timing of snowmelt. Because species closer in time to this unambiguous onset cue tended to be more strongly right-skewed, we can reasonably infer that this cue, or at least the phenological response of species to this cue, is less variable than the processes that lead to flower senescence and the end of bee foraging (e.g., frost events, precipitation, inherent lifespan/persistence, etc.). By extension, these findings suggest that later-season onset cues, or species’ phenological responses to them, are more variable than the early snowmelt cue.

Our finding of right-skewness in flowering phenology was broadly similar to previously published results, though we found that average flowering skewness in the present study (*g_1_* = 0.31) was less positively skewed than in a previous study in the same area (*g_1_* = 0.46; Thomson, 1980) and a prairie community (*g_1_* = 0.41, Rabinowitz et al., 1981). While this comparison is useful, we caution against over-interpretation due to differences between the studies such as duration of monitoring and size of sampling plots. Turning to insects, we lack other studies focused specifically on phenological skewness in other insect groups, but individual abundance time-series indicate that right-skewness may also be found in univoltine butterflies (Dennis et al., 2017; Scott et al., 1987; Zonneveld, 1991), flies (Haab et al., 2019; Judd et al., 1991), and hemipterans (Gamarra et al., 2020; Ma et al., 2008). Comparisons with multi-voltine species in areas with longer growing seasons are difficult, and further research is needed to compare uni- and multi-modal phenological distributions, especially as climate change creates opportunities for additional generations in some insects (Dyck et al., 2015; Hodgson et al., 2011). Given the apparent prevalence of skewness in phenological distributions, we encourage researchers to use modeling methods that are designed to capture asymmetry (Belitz et al., 2020). We advise caution when closely comparing skewness values between bees and flowers because there is inherently more uncertainty in the bee dataset due to the methodological challenges of tracking wild insects.

The consequences of variable skewness in flower and pollinator distributions for phenological match/mismatch in natural populations are not well understood and require further study. When considering simulated pair-wise interactions, the skewness of phenological distributions alone has the potential to cause up to 14% loss in overlap in the species that we studied (Fig. 4). We note that this analysis encompassed only phenological differences, and in reality there are other barriers to pollination such as specialization or morphological limitations to pollination. It is also important to consider that loss of overlap does not necessarily translate to fitness losses, as pollen limitation is not ubiquitous (Knight et al., 2005) and many bees and flowering plants are generalists (Waser et al., 1996). Beyond pollination interactions, differences in skewness have the potential to affect other mutualistic interactions, predator-prey and host-parasite interactions, and to alter patterns of inter- and intra-specific competition within guilds. As both flowers and bees tend to be right-skewed, individuals may compete most strongly with their conspecifics in the early part of their activity. Future studies should examine how phenological skewness translates into fitness consequences through changes in inter- and intra-specific interactions throughout species’ active periods.

## Acknowledgments

M.S. was funded by the NSF Graduate Research Fellowship under Grant No. 1745048. Field work for this research was supported by NSF grants DEB 94-08382, IBN-98-14509, DEB-0238331, DEB 0922080, and DEB-1912006 to DWI, DEB-0922080 and DEB-1354104 to D.W.I, B.D.I, N.U., and R.E.I., EarthWatch and its Research Corps, and funds from NC State University to R.E.I. We thank the many research assistants who have aided in the collection of the long-term bee and flower data, and the RMBL and private land owners for access to field sites. This research was conducted on the ancestral land of the Tabeguache band of the Ute people. Any opinions, findings, conclusions, or recommendations expressed in this material are those of the authors and do not necessarily reflect the views of the funding agencies.

## Notes

### Competing Interest Statement

The authors have declared no competing interest.

## References

Azzalini, A. (2020). The R package sn: The skew-normal and related distributions, such as the skew-t. CRAN. http://azzalini.stat.unipd.it/SN

Belitz, M. W., Larsen, E. A., Ries, L., & Guralnick, R. P. (2020). The accuracy of phenology estimators for use with sparsely sampled presence-only observations. Methods in Ecology and Evolution, 11(10), 1273–1285. https://doi.org/10.1111/2041-210X.13448

Blionis, G. J., Halley, J. M., & Vokou, D. (2001). Flowering phenology of *Campanula* on Mt Olympos, Greece. Ecography, 24(6), 696–706. https://doi.org/10.1111/j.1600-0587.2001.tb00531.x

CaraDonna, P. J., Iler, A. M., & Inouye, D. W. (2014). Shifts in flowering phenology reshape a subalpine plant community. Proceedings of the National Academy of Sciences, 111(13), 4916–4921. https://doi.org/10.1073/pnas.1323073111

Carter, S. K., Saenz, D., & Rudolf, V. H. W. (2018). Shifts in phenological distributions reshape interaction potential in natural communities. Ecology Letters, 21(8), 1143–1151. https://doi.org/10.1111/ele.13081

D’Agostino, R. B. (1970). Transformation to Normality of the Null Distribution of g1. Biometrika, 57(3), 679–681. https://doi.org/10.2307/2334794

Dennis, E. B., Morgan, B. J. T., Brereton, T. M., Roy, D. B., & Fox, R. (2017). Using citizen science butterfly counts to predict species population trends. Conservation Biology, 31(6), 1350–1361. https://doi.org/10.1111/cobi.12956

Dyck, H. Van, Bonte, D., Puls, R., Gotthard, K., & Maes, D. (2015). The lost generation hypothesis : could climate change drive ectotherms into a developmental trap? Oikos, 124, 54–61. https://doi.org/10.1111/oik.02066

Forrest, J., & Thomson, J. D. (2010). Consequences of variation in flowering time within and among individuals of *Mertensia fusiformis* (Boraginaceae), an early spring wildflower. American Journal of Botany, 97(1), 38–48. https://doi.org/10.3732/ajb.0900083

Gamarra, H., Sporleder, M., Carhuapoma, P., Kroschel, J., & Kreuze, J. (2020). A temperaturedependent phenology model for the greenhouse whitefly *Trialeurodes vaporariorum* (Hemiptera: Aleyrodidae). Virus Research, 289(198107). https://doi.org/10.1016/j.virusres.2020.198107

Gezon, Z. J., Wyman, E. S., Ascher, J. S., Inouye, D. W., & Irwin, R. E. (2015). The effect of repeated, lethal sampling on wild bee abundance and diversity. Methods in Ecology and Evolution, 6, 1044–1054. https://doi.org/10.1111/2041-210X.12375

Haab, K. A., McKnight, T. A., & McKnight, K. B. (2019). Phenology and ethology of adult *Lasiopogon slossonae* Cole and Wilcox robber flies (Diptera: Asilidae) in a New York riparian habitat. Proceedings of the Entomological Society of Washington, 121(4), 594–615. https://doi.org/10.4289/0013-8797.121.4.594

Hodgson, J., Thomas, C., Olver, T., Anderson, B., Brereton, T., & Crone, E. (2011). Predicting insect phenology across space and time. Global Change Biology, 17(3), 1289–1300. https://doi.org/10.1111/j.1365-2486.2010.02308.x

Inman, H. F., & Bradley, E. L. (1989). The overlapping coefficient as a measure of agreement between probability distributions and point estimation of the overlap of two normal densities. Communications in Statistics - Theory and Methods, 18(10), 3851–3874. https://doi.org/10.1080/03610928908830127

Inouye, B. D., Ehrlén, J., & Underwood, N. (2019). Phenology as a process rather than an event: from individual reaction norms to community metrics. Ecological Monographs, 89(2), 1–15. https://doi.org/10.1002/ecm.1352

Inouye, D. W. (2008). Effects of climate change on phenology, frost damage, and floral abundance of montane wildflowers. Ecology, 89(2), 353–362. https://doi.org/10.1890/06-2128.1

Judd, G. J. R., Whitfield, G. H., & Maw, H. E. L. (1991). Temperature-dependent development and phenology of pepper maggots (Diptera: Tephritidae) associated with pepper and horsenettle. Environmental Entomology, 20(6), 22–29. https://doi.org/10.1093/ee/20.1.22

Knight, T. M., Steets, J. A., Vamosi, J. C., Mazer, S. J., Burd, M., Campbell, D. R., Dudash, M. R., Johnston, M. O., Mitchell, R. J., & Ashman, T. L. (2005). Pollen limitation of plant reproduction: Pattern and process. Annual Review of Ecology, Evolution, and Systematics, 36, 467–497. https://doi.org/10.1146/annurev.ecolsys.36.102403.115320

Komsta, L., & Novomestky, F. (2015). Moments, cumulants, skewness, kurtosis and related tests. R package.

Ma, Z., & Bechinski, E. J. (2008). Developmental and phenological modeling of Russian wheat aphid (Hemiptera: Aphididae). Annals of the Entomological Society of America, 101(2), 351–361. https://doi.org/10.1603/0013-8746(2008)101[351:DAPMOR]2.0.CO;2

Ogilvie, J. E., Griffin, S. R., Gezon, Z. J., Inouye, B. D., Underwood, N., Inouye, D. W., & Irwin, R. E. (2017). Interannual bumble bee abundance is driven by indirect climate effects on floral resource phenology. Ecology Letters, 20, 1507–1515. https://doi.org/10.1111/ele.12854

Rabinowitz, D., Rapp, J. K., Sork, V. L., Rathcke, B. J., Reese, G. A., Weaver, J. C., Rapp, J. K., Reese, G. A., & Weaver, J. A. N. C. (1981). Phenological properties of wind- and insect-pollinated prairie plants. Ecology, 62(1), 49–56. https://doi.org/10.2307/1936667

Rathcke, B., & Lacey, E. P. (1985). Phenological patterns of terrestrial plants. Annual Review of Ecology and Systematics, 16, 179–214. https://doi.org/10.1146/annurev.es.16.110185.001143

Schmitt, J. (1983). Individual flowering phenology, plant size, and reproductive success in *Linanthus androsaceus*, a California annual. Oecologia, 59(1), 135–140. https://doi.org/10.1007/BF00388084

Scott, J. A., & Epstein, M. E. (1987). Factors affecting phenology in a temperate insect community. The American Midland Naturalist, 117(1), 103–118. https://doi.org/10.2307/2425712

Stemkovski, M., Pearse, W. D., Griffin, S. R., Pardee, G. L., Gibbs, J., Griswold, T., Neff, J. L., Oram, R., Rightmyer, M. G., Sheffield, C. S., Wright, K., Inouye, B. D., Inouye, D. W., & Irwin, R. E. (2020). Bee phenology is predicted by climatic variation and functional traits. Ecology Letters, 23(11), 1589–1598. https://doi.org/10.1111/ele.13583

Thomson, J. D. (1980). Skewed flowering distributions and pollinator attraction. Ecology, 61(3), 572–579. https://doi.org/10.2307/1937423

Visser, M. E., te Marvelde, L., & Lof, M. E. (2012). Adaptive phenological mismatches of birds and their food in a warming world. Journal of Ornithology, 153(Suppl 1), S75–S84. https://doi.org/10.1007/s10336-011-0770-6

Waser, N. M., Chittka, L., Price, M. V., Williams, N. M., & Ollerton, J. (1996). Generalization in pollination systems, and why it matters. Ecology, 77(4), 1043–1060. https://doi.org/10.2307/2265575

Zonneveld, C. (1991). Estimating death rates from transect counts. Ecological Entomology, 16(1), 115–121. https://doi.org/10.1111/j.1365-2311.1991.tb00198.x

